# Typical retinotopic locations impact the time course of object coding

**DOI:** 10.1101/177493

**Authors:** Daniel Kaiser, Merle M. Moeskops, Radoslaw M. Cichy

## Abstract

In everyday visual environments, objects are non-uniformly distributed across visual space. Many objects preferentially occupy particular retinotopic locations: for example, lamps more often fall into the upper visual field, whereas carpets more often fall into the lower visual field. The long-term experience with natural environments prompts the hypothesis that the visual system is tuned to such retinotopic object locations. A key prediction is that typically positioned objects should be coded more efficiently. To test this prediction, we recorded electroencephalography (EEG) while participants viewed briefly presented objects appearing in their typical locations (e.g., an airplane in the upper visual field) or in atypical locations (e.g., an airplane in the lower visual field). Multivariate pattern analysis applied to the EEG data revealed that object classification depended on positional regularities: Objects were classified more accurately when positioned typically, rather than atypically, already at 140 ms, suggesting that relatively early stages of object processing are tuned to typical retinotopic locations. Our results confirm the prediction that long-term experience with objects occurring at specific locations leads to enhanced perceptual processing when these objects appear in their typical locations. This may indicate a neural mechanism for efficient natural scene processing, where a large number of typically positioned objects needs to be processed.

## 1 Introduction

Visual objects are enclosed entities that can in principle be moved around freely. However, in everyday environments object positions are often quite constrained. For instance, consider the predictability in the locations of objects in a living room: The sofa is facing the TV, a table is in between the two, a lamp hangs from the ceiling, whereas carpets lie on the floor. This example illustrates that the object content of natural scenes is organized in repeatedly occurring positional structures (Bar, 2000; Chun, 2002). Many previous studies have investigated how inter-object relationships in these positional structures (e.g., lamps appearing above tables) impact behavioral performance and neural processing (Biederman, Mezzanotte, & Rabinowitz, 1982; Kaiser, Stein, & Peelen, 2014; Oliva & Torralba, 2007; Wolfe, Võ, Evans, & Greene, 2011). However, positional object structures often also imply that individual objects are associated with particular locations in space (e.g., lamps appearing in the upper part of a scene). It has recently been proposed that the visual system is tuned to these regularities (Kaiser & Haselhuhn, 2017; Kravitz, Vinson, & Baker, 2008), which could facilitate neural processing for individual objects appearing in retinotopic locations that correspond to their typical real-world locations.

Such location-specific variations in object coding are suggested by previous results that indicate the co-representation of object identity and location information in visual cortex: (1) cortical responses depend on the position of the object in the visual field (Hemond, Kanwisher, & Op de Beeck, 2007; Hasson, Levy, Behrmann, Hendler, & Malach, 2002), (2) object selective cortex contains information about both an object’s identity and its location (Cichy, Chen, & Haynes, 2011; Golomb & Kanwisher, 2011; Hong, Yamins, Majaj, & DiCarlo, 2017; Kravitz, Kriegeskorte, & Baker, 2010; Schwarzlose, Swisher, Dang, & Kanwisher, 2008), and (3) information about object identity and location emerge at similar time points in visual processing (Isik, Meyers, Leibo, & Poggio, 2014; Carlson, Hogendoorn, Kanai, Mesik, & Turret, 2011).

The link between identity and location information in object processing creates the possibility that the two properties interact. In everyday environments, the visual system is repeatedly faced with positional structures, where individual object positions are highly predictable. Under typical viewing conditions, and unless directly fixated, objects appearing in the upper part of scenes also more often occupy locations in the upper visual field, while objects appearing in the lower part of scenes are repeatedly encountered in the lower visual field. Through repeated exposure, retinotopic object-coding mechanisms could get tuned to typical object locations in the visual field. Thus, over time, neural channels are shaped to represent typical object-location conjunctions in a maximally efficient way. These efficient location-specific object representations would enhance the processing of an object when it appears in its typical locations within a scene – and within the visual field. Evidence for such a processing enhancement has been found in the domain of person perception, where typical configurations impact cortical responses to individual face and body parts (Chan, Kravitz, Truong, Arizpe, & Baker, 2010; de Haas et al., 2016; Henriksson, Mur, & Kriegeskorte, 2015). For example, in face-selective visual cortex, response patterns are better discriminable for typically, as compared to atypically, positioned face parts (de Haas et al., 2016), revealing visual processing channels that are tuned to the spatial regularities in the face.

Here, we test the prediction that the positional regularities contained in natural scenes can similarly facilitate the processing of everyday objects appearing in their typical retinotopic locations. Participants viewed objects associated with upper and lower visual field locations (e.g., a lamp or a carpet) (Figure 1A) while we recorded electroencephalography (EEG). We used multivariate classification on the EEG data (Contini, Wardle, & Carlson, 2017) to track the time course of object coding with high temporal precision. Analyses revealed that after 140ms the visual processing of an object is affected by its typical location in the visual field: Objects appearing in their typical locations (e.g., a lamp in the upper visual field and a carpet in the lower visual field) could be decoded more successfully than objects appearing in atypical locations (e.g., a carpet in the upper visual field and a lamp in the lower visual field). These results suggest that early stages of visual processing are tuned to the positional object structure of real-world scenes.

**Figure 1.**
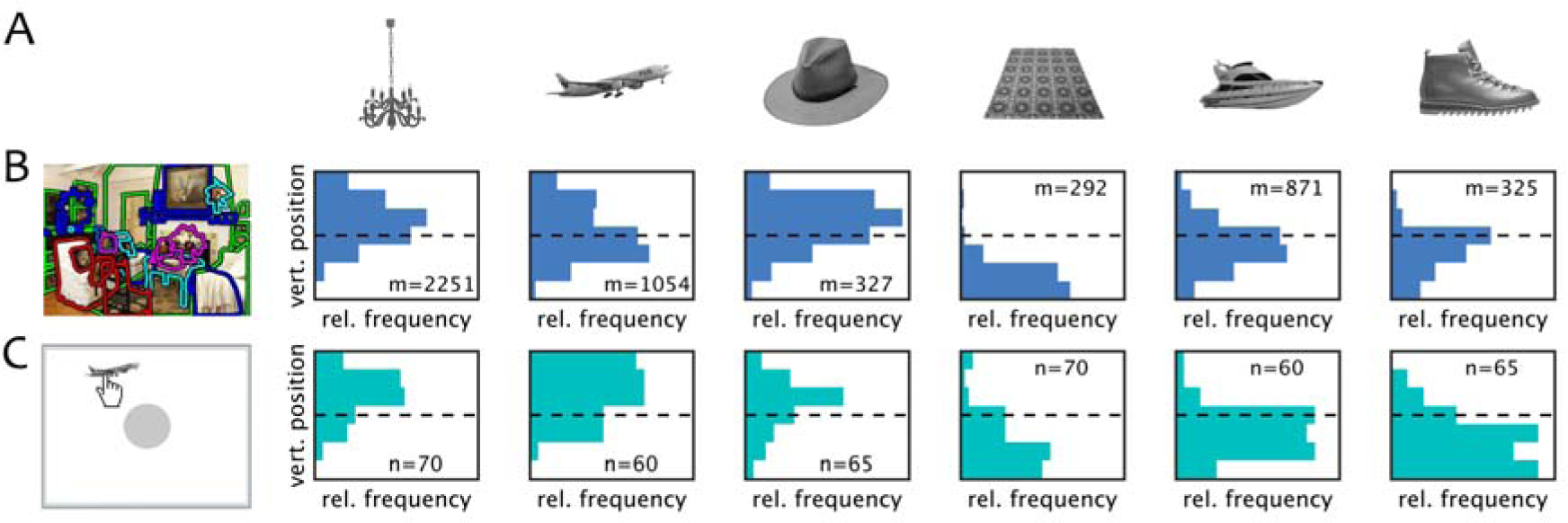
A) Example Stimuli. Ten different exemplar images of six objects each (here, one example image for each object shown) were used as stimuli, of which three objects were associated with the upper visual field (lamp, airplane, hat) and three were associated with the lower visual field (carpet, boat, shoe). B) To validate our assessment of visual field associations, we automatically extracted the positions for each object in a large set of labelled scene photographs taken from the LabelMe scene database (Russell et al., 2008). For each scene, we determined the relative position of the object along the vertical axis, and plotted the distribution across 7 bins (*m*: number of scenes for each object). C) Additionally, we asked a group of participants to indicate for each object the position that it typically occupies in the visual field by dragging the object to the desired location. We then computed the distribution of relative locations along the vertical axis of the screen, split into 7 bins (n: number of participants that indicated the typical location for each object). Both measures confirmed the spatial priors associated with the six objects.

## 2 Materials and Methods

### 2.1 Participants

Thirty-four healthy adults (mean age 26.4 years, *SD* = 5.4; 23 female) completed the experiment. The sample size was set a-priori, based on considerations regarding statistical power: A sample size of 34 is needed for detecting a simple effect with a medium effect size of *d* = 0.5 with a probability of more than 80%. All participants had normal or corrected-to-normal vision, provided informed consent and received monetary reimbursement or course credits for their participation. All procedures were approved by the ethical committee of the Department of Education and Psychology of the Freie Universität Berlin and were in accordance with the Declaration of Helsinki.

### 2.2 Stimuli

The stimulus set consisted of greyscale images of six objects associated with typical visual field locations, of which three were associated with upper visual field locations (lamp, airplane, and hat) and three were associated with lower visual field locations (carpet, boat, and shoe). For each object, ten exemplars were used (see Figure 1A for stimulus examples). To avoid a confounding of visual field associations and other, conceptual stimulus attributes, the objects were matched for their categorical content (two furniture items, two transportation items, and two clothing items), so that high-level properties like the objects’ size, manipulability and semantic associations were comparable across upper and lower visual field objects. To control for low-level visual factors, all stimulus images were gray-scaled and matched for overall luminance (using the SHINE toolbox; Willenbockel et al., 2010). Additionally, as a measure of low-level image similarity, pixel-wise similarity was computed between all 60 images (i.e., 6 objects, 10 exemplars each), in a pairwise fashion: Pixel intensity values were correlated between all same visual field objects (e.g., lamps and airplanes) and between all different visual field objects (e.g., lamps and boats). No difference between all possible (600) within-field and all possible (900) between-field correlations emerged, *t*(1498)=0.50, *p*=0.62, indicating that objects with identical typical locations were no more similar in their low-level visual properties than objects with different typical locations.

To ensure that the six objects could be reliably linked to a specific location, we validated the association of the six objects with a specific part of the visual field in two ways. First, we assessed the typical spatial distribution of each object in natural scenes, assuming that natural scene photographs represent a snapshot of the visual field roughly approximating natural viewing conditions. Hence, the distribution of the objects in the scene image should be similar to their distribution across the visual field. To objectively measure the typical position of each object within a scene, we queried a huge number of scene photographs (>100,000) from the LabelMe toolbox, where human observers annotated single objects by drawing labelled polygons (Russell, Torralba, Murphy, & Freeman, 2008). For all scenes containing a specific object we computed the mean pixel coordinate of the area labeled as belonging to the object and then averaged these positions across scenes. The resulting “typical” object locations showed that, as expected, the upper visual field objects were associated with locations (*y*: vertical coordinate from bottom (0) to top (1) of the scene) in the upper parts of scenes (lamp: *y* = 0.61, *SD* = 0.17; airplane: *y* = 0.52, *SD* = 0.20; hat: *y* = 0.60, *SD* = 0.18), while lower visual field objects were associated with locations in the lower part of scenes (carpet: *y* = 0.17, *SD* = 0.13; boat: *y* = 0.42, *SD* = 0.18; shoe: *y* = 0.37, *SD* = 0.17). The typical location in scenes differed significantly between objects associated with the upper and lower visual field, t > 11.4, *p* < .001, for all pairwise comparisons. Figure 1B shows the distribution of object locations along the vertical axis of the scenes, split into 7 bins.

Second, we sought to demonstrate a correspondence between this automated, scene-based measure and people’s explicit associations of the objects with particular locations in space. We thus asked a set of participants (n = 70 for lamp/carpet, n = 60 for airplane/boat, n = 65 for hat/shoe; including the participants of the current study, after the completion of the EEG experiment) to indicate the typical locations in which they expect to see each of the six objects. In this task, participants were asked to drag the image of a single exemplar of each object to its typical location on a computer screen (imagining that the computer screen represents their field of view in a natural scene). The central part of the screen – where the object initially appeared – was blocked (indicated by a grey circle), so that participants (1) could not place the object in a central location of the screen, and (2) had to move the object before proceeding. As expected, participants more often chose upper screen positions (y: vertical coordinate from bottom (0) to top (1) of the screen) for the upper visual field objects (lamp: *y* = 0.65, *SD* = 0.19; airplane: *y* = 0.67, *SD* = 0.20; hat: *y* = 0.57, *SD* = 0.20), and lower screen positions for the lower visual field objects (carpet: *y* = 0.29, *SD* = 0.22; boat: *y* = 0.36, *SD* = 0.18; shoe: *y* = 0.30, *SD* = 0.18). The vertical locations chosen by the participants differed significantly between objects associated with the upper and lower visual field, *t* > 6.04, *p* < .001, for all pairwise comparisons. Figure 1C shows the distribution of vertical object locations on the screen, split into 7 bins. The scene-based measure and participants’ explicit assessment thus provided converging evidence for the association of the objects with specific spatial locations.

### 2.3 Experimental Design

To test whether objects are processed differently when presented in typical and atypical locations within the visual field, the objects were presented in the upper or lower visual field (Figure 2A). On every trial, one object exemplar was presented in one of the two locations for 150ms, followed by a variable inter-trial interval (randomly jittered, from 1250ms to 1750ms). The brief presentation time was chosen to not give participants enough time to move their eyes towards the objects. Stimuli were presented at 3.25° vertical eccentricity and subtended a visual angle of maximally 3° in horizontal and vertical axes. Stimulus presentation was controlled using the Psychtoolbox (Brainard, 1997). Participants were asked to detect one-back repetitions on an object level (e.g., two different airplanes in direct succession; see Figure 2A). Repetitions occurred on 13% of the trials and equally often for typically and atypically positioned repetition targets and for the top and bottom locations. Participants performed accurately on this task (mean accuracy 96%, *SD* = 2%), with no difference in accuracy between typically and atypically positioned objects, *t*(33) = 0.87, *p* = .391. One-back repetition trials (i.e., the “second” trial containing the repetition) were removed from all EEG analyses. The whole experiment consisted of 1656 trials: in 1440 trials, each object exemplar was shown 12 times in each location (i.e., 120 repetitions per object and location), while the remaining 216 trials were one-back repetition trials. The experiment was split into 8 runs, and participants could take breaks between the runs. Twelve participants completed an extended experimental session with 2760 trials (including 360 repetition trials), which additionally contained the same conditions at large eccentricities in half of the trials; these additional data are not reported here. The 2400 non-repetition trials consisted of 10 repetitions of each object exemplar in each of the four locations (i.e., 100 repetitions per object and location). The extended experiment was split into 12 runs.

**Figure 2.**
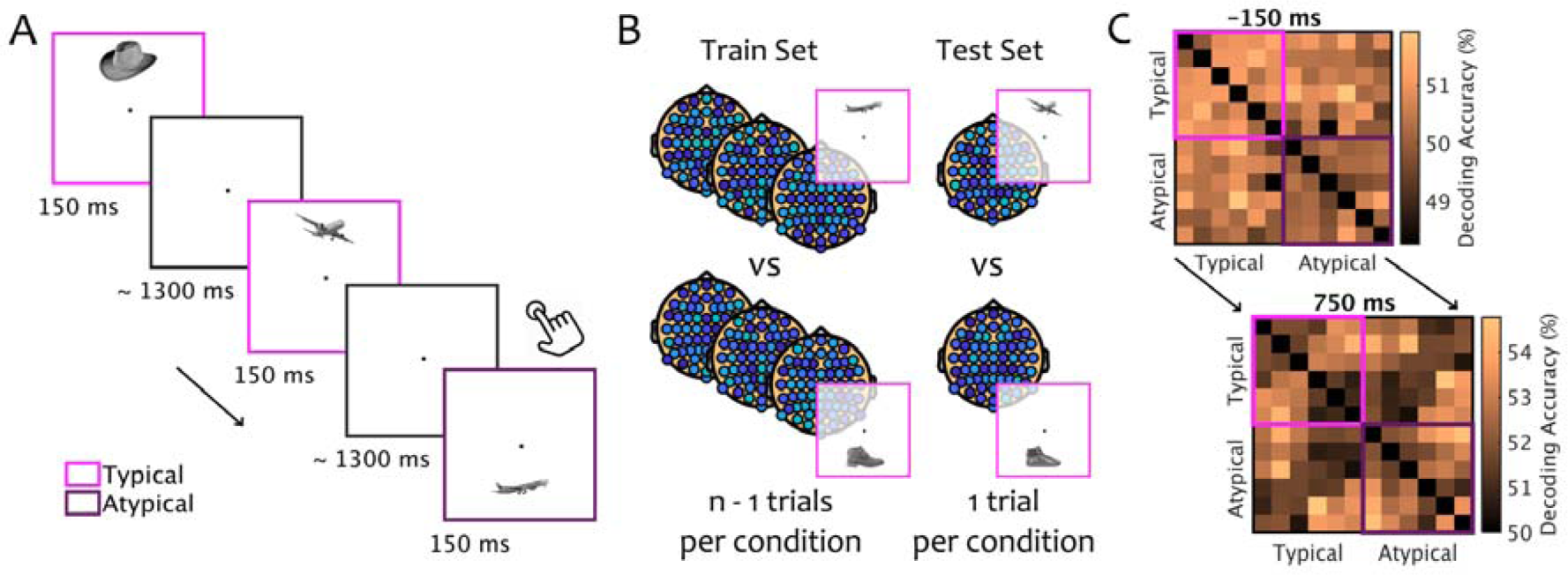
Paradigm and Classification Logic. A) Stimuli were presented for 150 ms in upper or lower visual field locations, corresponding to an object’s typical or an atypical location. Participants were instructed to detect occasional one-back repetitions on an object-level (e.g., two airplanes in a row; irrespective of the stimulus location) by pressing a button. Colors indicating the two regularity conditions are shown for illustrative purposes only. B) Multivariate classification was performed on response patterns across all electrodes, separately for each pairwise combination of objects (exemplified here by airplane and shoe in regular locations). The data was split into two sets: a training set consisting of all (but one) trials for each object and a testing set consisting of the two left-out trials. LDA classifiers were repeatedly trained and tested until every trial was left out once and accuracy was averaged across these repetitions. C) The pairwise classification analysis was repeated for each 10 ms time bin, resulting in a 12-by-12 matrix of pairwise classification accuracies (with an empty diagonal) at every time point. To determine differences between typically and atypically positioned objects, pairwise comparisons within the typically placed objects (pink rectangle, upper left) and within the atypically placed objects (purple rectangle, lower right) were averaged and compared (Figure 3).

### 2.4 EEG recording and preprocessing

The EEG was recorded using an EASYCAP 64-channel system and a Brainvision actiCHamp amplifier. The 64 electrodes were arranged in accordance with the standard 10-10 system. The data was recorded at a sampling rate of 1000 Hz and filtered online between 0.5 and 70 Hz. For one participant, due to a technical problem, only data from 32 electrodes was recorded. All electrodes were referenced online to the Fz electrode. Offline preprocessing was performed in MATLAB, using the FieldTrip toolbox (Oostenveld, Fries, Maris, & Schoffelen, 2011). The continuous EEG data was epoched into trials ranging from 150ms before stimulus onset to 750ms after stimulus onset. Trials containing movement-related artefacts were visually identified and excluded from all analyses. Blink and eye movement artifacts were identified and removed using Independent Components Analysis (ICA) and visual inspection of the resulting components. To increase the signal-to-noise ratio of the classification analyses (Carlson, Tovar, Alink, & Kriegeskorte, 2013), the data was downsampled to 100Hz.

### 2.5 EEG classification procedure

Multivariate classification analyses were carried out in MATLAB using the CoSMoMVPA toolbox (Oosterhof, Connolly, & Haxby, 2016). Classification was performed separately for each 10ms time bin, resulting in classification time courses with 10 ms resolution. The analysis was performed pairwise, for all possible combinations of the six objects appearing in the two locations. Linear discriminant analysis (LDA) classifiers were always trained and tested on data from two conditions (e.g., an airplane in the upper visual field versus a carpet in the lower visual field), using a leave-one-out partitioning scheme (Figure 2B). The training set consisted of all but one trials for each of the two conditions, while one trial for each of the two conditions was held back and used for classifier testing. This procedure was repeated until every trial was left out once. Classifier performance was averaged across these repetitions. The pairwise decoding analysis resulted in 12-by-12 matrix of decoding accuracies at each time point (reflecting all comparisons between the six objects appearing in the two locations) (Figure 2C).

### 2.6 Overall classification dynamics

To assess the overall classification dynamics over time, we computed the general discriminability of the twelve different conditions. All pairwise classification accuracies were averaged, revealing a time course of object decoding independently of the positional regularities. This time course of overall classification accuracy was used to define time points of interest at the peaks of the classification time series, where classification performance was particularly pronounced. Using a “region of interest” logic frequently applied in fMRI analyses (Poldrack, 2007), we used these peaks as “time points of interest” to increase the detection power of subsequent analyses.

### 2.7 Object classification in typical and atypical locations

To determine an effect of positional regularity on object decoding, we compared performance when classifying among typically positioned objects versus among atypically positioned objects. Pairwise classification accuracies were averaged for all comparisons between typically positioned objects (e.g., an airplane in the upper visual field versus a shoe in the lower visual field) and for all comparisons between atypically positioned objects (e.g., a shoe in the upper visual field versus an airplane in the lower visual field) (Figure 2C). Subsequently, the two resulting classification time series (for typically and atypically positioned objects) were compared. To increase the statistical power of this comparison, we specifically focused on the effect of positional regularity at the peaks in overall classification.

### 2.8 Sensor-space searchlight analysis

To investigate which sensors contributed most to the observed effects, we performed a sensor-space searchlight analysis. For this analysis, the pairwise classification procedure was repeated for neighborhoods of seven adjacent electrodes around each individual electrode; the resulting classification accuracy was then mapped onto a scalp representation. This procedure allowed us to infer the approximate spatial distribution of classification differences between typically and atypically positioned objects. As for one participant only data from 32 electrodes was available, this participant was not included in the searchlight analysis.

### 2.9 Statistical testing

Statistical testing was done across participants. To identify significant effects across time we used a threshold-free cluster-enhancement procedure (Smith & Nichols, 2009) with default parameters. Multiple comparison correction was based on a sign-permutation test (with null distributions created from 10,000 bootstrapping iterations) as implemented in CoSMoMVPA (Oosterhof et al., 2016). The resulting statistical maps were thresholded at *Z* > 1.96 (i.e., *p* < .05). The same procedure was employed for identifying significant sites across electrodes in the sensor-space searchlight analysis. For assessing the significance of effects at the overall classification peaks, repeated-measures ANOVAs and paired t-tests were performed. For these tests, we also computed effect size measures: partial *η*^2^ for ANOVAs and Cohen’s d for t-tests. Additionally, for t-tests, scaled-information Bayes factors (BF) were calculated as an additional measure of evidential value (Rouder, Speckman, Sun, Morey, & Iverson, 2009), where Bayes factors >10 can be considered strong evidence for an effect.

## 3 Results

### 3.1 Temporal dynamics of pairwise object classification

In a first step, we characterized the overall response dynamics observed in the pairwise classification analysis, which allowed us to restrict subsequent analyses to time points where classification performance was particularly pronounced. For this, we computed an overall measure of pairwise classification by averaging across all unique off-diagonal elements of the pairwise classification matrices (Figure 2C), resulting in a single classification time series. This analysis revealed robust above-chance classification starting from 70 ms after stimulus onset and prominently peaking at 140 ms and 220 ms (Figure 3A). These two clear peaks in the classification time series were used as time points of interest for subsequent analyses, as we reasoned that differences between typically and atypically positioned objects would be most pronounced at time points at which objects were most discriminable.

**Figure 3.**
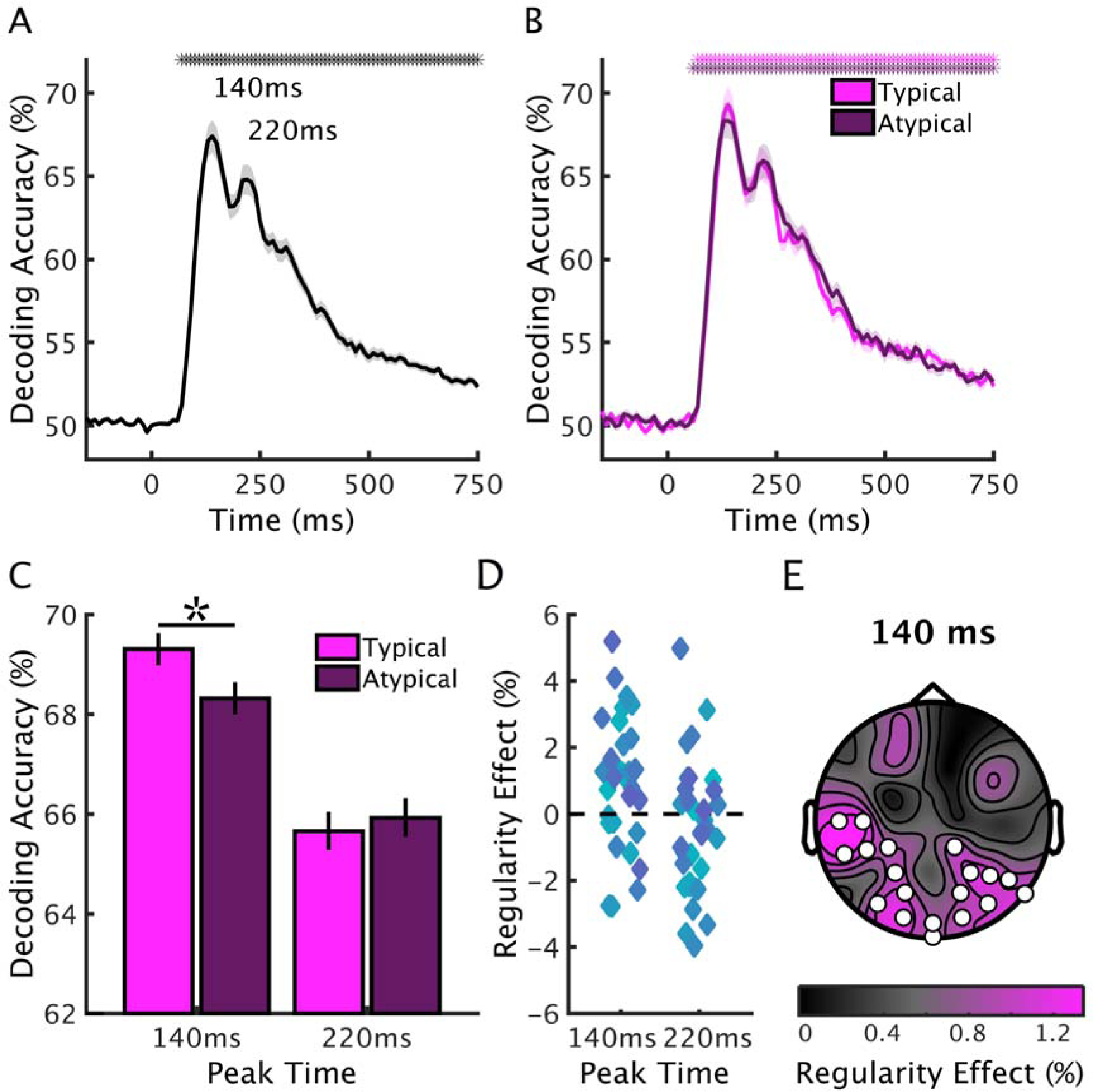
Classification Results. A) Overall pairwise classification performance was computed by averaging all pairwise decoding time series, revealing significant decoding accuracy starting at 70 ms after stimulus onset and peaking at 140 ms and 220 ms. Asterisks above data curves indicate above-chance classification (*p* < .05, corrected for multiple comparisons). The shaded margin reflects standard errors of the mean. B) Classification time series for typically and atypically positioned objects were computed by averaging all pairwise classification time series comparing typically and atypically positioned objects, respectively. Classification of typically and atypically positioned pairs showed comparable temporal dynamics, both peaking at the time points identified in the overall decoding. Asterisks indicate above-chance classification (*p* < .05, corrected for multiple comparisons). The shaded margins reflect standard errors of the mean. C) At the first decoding peak (140 ms), but not the second peak (220 ms), classification was more accurate for typically than for atypically positioned objects. The asterisk indicates a significant difference (*p* < .05). Error bars reflect standard errors of the difference. D) Scatterplots showing the regularity effects across participants, for both peak times. E) A sensor-space classification searchlight revealed that the regularity effect (difference between the classification of typically and atypically positioned objects) at the 140 ms peak is most pronounced in occipital and temporal electrodes. Circles indicate electrodes exhibiting a significant regularity effect (*p* < .05, corrected for multiple comparisons).

### 3.2 Classification of objects when positioned typically and atypically

To test whether neural representations differ for typically and atypically positioned objects, we compared classification performance for all typically and all atypically positioned objects. We averaged all pairwise classification time courses that corresponded to comparisons within regular pairs (e.g., an airplane in the upper visual field versus a boat in the lower visual field) and comparisons within irregular pairs (e.g., an airplane in the lower visual field versus a boat in the upper visual field) (Figure 3B). The classification time series for typically and atypically positioned objects showed a similar temporal structure and both replicated the peak structure observed in the overall decoding, allowing for a meaningful comparison between typically and atypically positioned objects at the classification peaks. We thus restricted statistical comparisons to two time points of interest: the peak times observed in the overall decoding (140 ms and 220 ms). For the early peak at 140ms, we found higher classification accuracy for typically than for atypically positioned objects, *t*(33) = 3.04, *p* = .005, *d* = 0.52, *BF* = 12.06, while for the later peak at 220ms, no such difference emerged, *t*(33) = 0.69, *p* = .495, *d* = 0.12, *BF* = 3.37, interaction with peak time, *F*(1,33) = 7.44, *p* = .010, *η*^2^ = 0.18 (Figure 3C). This pattern of results suggests that earlier stages of object processing (as reflected in the decoding peak at 140 ms) benefit from typical object locations, while relatively later object representations (at 220 ms after stimulus onset) are not sensitive to positional regularities. This result was replicated in a bootstrapping analysis, where we used independent sub-groups of participants to define peak times and compute the regularity effects (see Supplementary Information, Fig. S1).

To estimate the spatial extent of the early regularity effect, we performed a searchlight analysis in sensor space. We repeatedly performed the pairwise classification analysis for neighborhoods of seven adjacent sensors, using only data from the early peak at 140 ms. To quantify the regularity benefit, we then computed the difference between all pairwise comparisons of typically positioned objects and all pairwise comparisons of atypically positioned objects at every sensor location. This analysis revealed a significant regularity effect in posterior and lateral electrodes (19 significant electrode sites) (Figure 3D). This result provides an indication of the tentative cortical source of the regularity effect (within the limits of the restricted spatial resolution of EEG measurements), suggesting that the enhanced classification for regularly positioned objects may originate from visual areas of the occipital and temporal cortices.

These results also provide a control for oculomotor influences on our findings. If differential eye-movement patterns were driving the enhanced decoding for typically positioned objects, the effect should be observed in frontal electrodes, too. The absence of a regularity benefit in the frontal electrodes thus provides evidence against eye-movement confounds. Two additional control analyses focused on spatiotemporal response patterns in selected frontal electrodes (see Supplementary Information, Fig. S3); these analyses confirmed the absence of an effect, further rendering eye-movement confounds unlikely.

### 3.3 Classification within and between locations

Our classification approach collapsed across pairwise comparisons within the same location and between different locations, so that classifiers could rely on information from both an object’s identity and its location. Therefore, the classification benefit for typically positioned objects could, in principle, also emerge from a more thorough coding of an object’s location. For example, classifiers – in principle – only need to predict the object’s location to correctly classify a lamp in the upper visual field versus a shoe in the lower visual field. By contrast, to correctly classify a lamp in the upper visual field versus a hat in the upper visual field, classifiers need to use information about the object. Note that both these comparisons contributed to the classification accuracy of regularly positioned objects in the previous analysis.

To test whether a regularity benefit emerges also in situations where classification solely depends on object information, we thus looked at the regularity effect for all comparisons between locations (e.g., an airplane in the upper visual field versus a shoe in the lower visual field) (Figure 4A) and all comparisons within location (e.g., an airplane in the upper visual field versus a hat in the upper visual field) (Figure 4B). If typical positioning enhances location information, a regularity effect should only be found when comparing objects between different locations, but not within a specific location (when location information is eliminated). Finding comparable effects in the within- and between-location comparisons would not support this view, indicating that typical positioning can also enhance processing when only object information is diagnostic.

**Figure 4.**
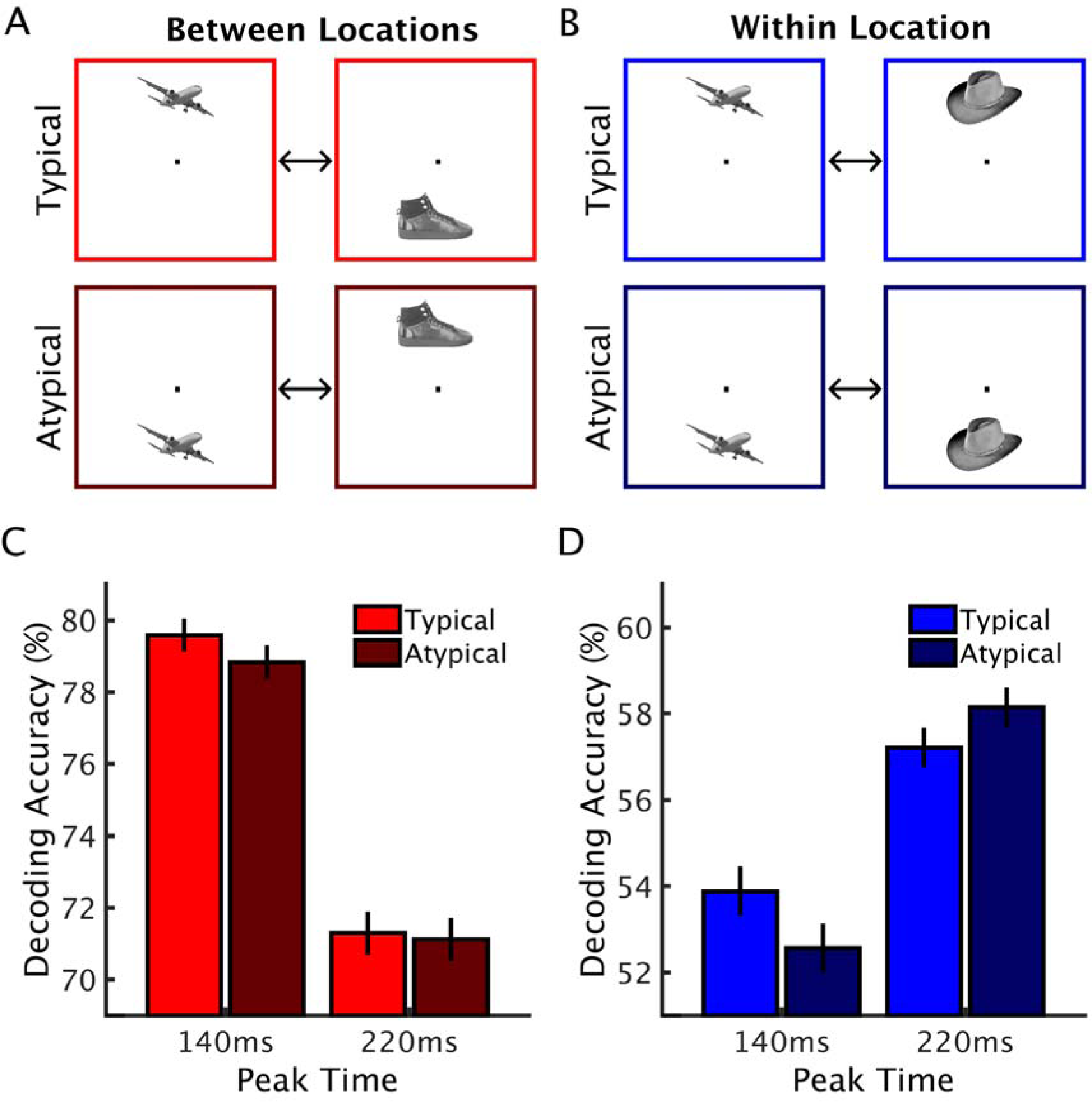
Between-Locations versus Within-Location Classification. We compared peak decoding for typically and atypically positioned objects separately for comparisons between different locations (e.g., an airplane in the upper visual field versus a shoe in the lower visual field) (A) and within the same location (e.g., an airplane in the upper visual field versus a hat in the upper visual field) (B). For both comparison types, we found a similar pattern (C, D) with a benefit for typically positioned objects at the 140 ms peak. Importantly, the patterns of results for between-locations and within-location classification were statistically indistinguishable (see Results). Error bars reflect standard errors.

We found a main effect of visual field comparison, *F*(1,33) = 365.70, *p* < .001, *η*^2^ = 0.91, with higher classification accuracies for classifying between locations (where the classifier can use the stimulus’ location) than within location (where the classifier has no location information available), and an interaction of the within-between comparison and peak latency, *F*(1,33) = 72.30, *p* < .001, *η*^2^ = 0.69, with a relatively more pronounced early peak when classifying between locations. Replicating our previous results, the analysis produced a significant peak X regularity interaction, *F*(1,33) = 9.83, *p* = .004, *η*^2^ = 0.23, with a regularity benefit at the 140 ms peak, *t*(33) = 3.15, *p* = .004, *d* = 0.54, BF = 15.52, but not the 220 ms peak, *t*(33) = 1.05, *p* = .301, *d* = 0.18, *BF* = 2.50. Crucially, this pattern of results did not depend on the type of classification (between locations versus within location), *F*(1,33) = 2.53, *p* = .121, *η*^2^ = 0.07. These results do not provide evidence for a boost of location information, thus suggesting that typical real-world locations enhance visual processing by boosting object identity information.

## 4 Discussion

### 4.1 Summary

Here, we demonstrate that positional regularities contained in real-world scenes impact brain responses to individual objects. Using multivariate classification of EEG data, we show that object coding across the visual field is affected by the typical real-world location of the object. When objects are presented in frequently experienced locations, EEG response patterns at 140 ms after stimulus onset are better discriminable than when the same objects are presented in atypical locations. This early regularity benefit yielded a robust effect size of *d* = 0.52, and was consistent across subsets of participants (see Supplementary Information). Interestingly, the advantage for typically positioned objects was equally pronounced for classification between locations and within the same location, suggesting that typical positioning boosts object identity information, rather than location information. Using a sensor-space searchlight analysis, we characterize the effect in sensor space, where its most prominent emergence in posterior sensors suggests that typically positioned objects gain an advantage during early perceptual processing.

### 4.2 Retinotopic priors as a consequence of natural scene structure

How does this retinotopically specific processing benefit emerge from the structure of natural environments? An intuitively appealing explanation lies in the interplay of typical within-scene object locations and gaze patterns during natural viewing. When objects repeatedly occupy specific locations within a scene, they may repeatedly fall into similar parts of the visual field, thereby shaping visual tuning properties. For instance, as lamps mostly appear in the “upper” part of a scene, fixations likely more often land below a lamp than above a lamp. Previous eye-tracking studies can provide some evidence for such a relationship between within-scene locations and retinotopic locations in limited experimental contexts: e.g., when walking along a corridor, lamps most often fall in the upper visual field (Turano et al., 2003). However, gaze patterns in real-world situations are complex and vary as a function of task context (Hayhoe & Ballard, 2005; Land, 2009), making it hard to establish a comprehensive characterization of this relationship. Future large-scale studies that investigate viewing patterns under naturalistic viewing conditions are thus required to empirically establish the connection between typical within-scene locations and typical retinotopic locations with high ecological validity. Without such studies, the association between typical within-scene locations and retinotopic priors – although intuitively compelling – needs to remain somewhat speculative.

### 4.3 Early stages of object coding are sensitive to typical locations

Our findings demonstrate that visual processing channels are preferentially tuned to specific objects appearing in specific retinotopic locations (Kaiser & Haselhuhn, 2017; Kravitz et al., 2008). Crucially, our EEG classification approach allowed us to pinpoint the latency of this regularity benefit: We demonstrate that object processing at 140 ms after stimulus onset is affected by positional regularity. The timing of the effect suggests that objects appearing in typical visual field locations gain an advantage during early, perceptual processing, rather than through top-down interactions (e.g., via long-range feedback from frontal areas); previous M/EEG studies have suggested that such feedback processes impact visual responses only at later stages, starting shortly before 200 ms (Bar et al., 2006; Fahrenfort, van Leeuwen, Olivers, & Hogendoorn, 2017). As opposed to the difference in early object processing, later representations (at 220 ms after stimulus onset) do not depend on the location of the object. This result concurs with the increasing location tolerance over the time course of object classification, peaking at around 180 ms (Isik et al., 2014), mirroring the increase in receptive field size along the visual stream (Kravitz, Saleem, Baker, Ungerleider, & Mishkin, 2013).

Can the sensitivity to positional regularity be interpreted as a processing advantage for typically positioned objects? Or do the results rather reflect a processing disadvantage for atypically positioned objects? Alternatively, there might be both an experience-based advantage for typically positioned objects, and a disadvantage for atypically positioned objects, caused by a lack of such experience? Our design cannot conclusively disambiguate these possibilities, as it relies on relative effects (i.e., comparing typical and atypical positioning). In the Supplementary Information, we however provide indirect evidence by analyzing individual-object effects: decoding of typically positioned objects increases when the object has a stronger location prior, while the decoding of atypically positioned objects does not vary as a function of the prior strength (Fig. S2). Although these results provide tentative evidence for a processing advantage for typically positioned objects (rather than a disadvantage for atypically positioned objects), future investigations need to test the different interpretations more explicitly, for example by including objects that are not associated with specific locations.

### 4.4 Visual versus categorical sources of the regularity benefit

What is the content of the location-specific object representations emerging at 140 ms? Peaks in the M/EEG decoding in this time range have been previously associated with visual category processing in object-selective cortex (Cichy, Pantazis, & Oliva, 2014, 2016; Carlson et al., 2013). Our searchlight analysis reveals the strongest regularity effect over lateral occipital and temporal electrode sites, suggesting that the effect originates from object-selective visual cortex. Whether processing differences in these object-selective regions reflect genuine category processing differences or whether they reflect differential coding of category-associated visual features is a debated question (Bracci, Ritchie, & op de Beeck, 2017; Peelen & Downing, 2017). While some data suggest that visual properties explain most of the variance in object-selective responses (e.g., Baldassi et al., 2013), a recent MEG decoding study has demonstrated category-selective, rather than visually (shape-) driven, responses from as early as 130 ms after stimulus onset (Kaiser, Azzalini, & Peelen, 2016). To determine if the regularity benefit observed here can be linked to differences in the processing of particular visual features or true categorical processing differences, future studies need to employ stimuli that vary more extensively in their visual characteristics.

### 4.5 Positional structures beyond person perception

Positional regularities have been studied in humans and non-human primates largely in the context of face and body perception, where parts are arranged in highly predictable configurations (e.g., the features of a human face). fMRI studies in humans have demonstrated that individual face and body parts are processed more efficiently when they appear in typical visual field locations (Chan et al., 2010; de Haas et al., 2016). Single-cell recordings in monkeys demonstrated that location biases can impact cortical responses to face parts as early as 100 ms after stimulus onset (Issa & DiCarlo, 2012), suggesting a benefit at early stages of perceptual processing.

Our results complement these findings by showing that such location-specific object processing is not restricted to the face/body domain: The inherent structure of natural scenes can similarly impact early processing (140 ms after stimulus onset) of object information across the visual field. Our findings thus highlight that location-specific tuning in object processing may form a general principle that shapes visual processing mechanisms for spatially predictable information. Future research could test whether regularity structures also affect other domains where the visual input consists of multiple parts that are constrained by spatial regularities. For example, through extensive experience with reading written text, the neural mechanisms for perceiving letters could get tuned to their typical spatial locations within words (Kaiser & Haselhuhn, 2017; Vinckier et al., 2007).

### 4.6 Positional structures in multiple and individual objects

Natural environments contain positional regularities on different levels, both on the levels of multiple (e.g., a lamps typically hang above tables) and individual objects (e.g., a lamp is typically in the upper visual field). Previous research has primarily focused on the latter: Recent behavioral studies have demonstrated that regularity structures in multi-object arrangement facilitate behavior in capacity-limited visual tasks (Gronau & Shachar, 2014; Kaiser et al., 2014; Kaiser, Stein, & Peelen, 2015; Stein, Kaiser, & Peelen, 2015), and neuroimaging studies demonstrated that they enable the brain to integrate information across objects that appear in frequently experienced arrangements (Baeck, Wagemans, & Op de Beeck, 2013; Kaiser & Peelen, 2018; Kaiser et al., 2014).

Here, we provide the first evidence that typical regularity structures also impact the neural representation of individual objects. Our finding thus raises the question whether the previously reported regularity effects in multi-object perception can be reduced to the effects of typical individual object location. On a behavioral level, some previous studies oppose this notion by demonstrating that the benefits of multi-object regularities cannot be explained by the relative location of the constituent objects (Kaiser et al., 2014, 2015; Stein et al., 2015). Although these results suggest that positional regularities in multi-object and single-object processing offer complementary benefits, further research is needed. Future studies will need to systematically manipulate positional structures on different levels (from individual objects to multi-object arrangements) to explore how regularities on multiple levels interact on a neural level.

### 4.7 Object processing in the context of natural scenes

A major challenge for the visual system when processing natural scenes is the large number of individual objects they contain. Surprisingly, however, objects can be rapidly decoded from MEG activity patterns, even when embedded in complex scenes (Brandman & Peelen, 2017; Kaiser, Oosterhof, & Peelen, 2016). Previous electrophysiological studies have investigated how congruent scene context can facilitate cortical processing of an object. Several studies have shown that semantic consistencies (i.e., whether the object is associated with the scene) affect EEG waveforms starting from around 300 ms (Ganis & Kutas, 2003; Mudrik, Lamy, & Deouell, 2010; Mudrik, Shalgi, Lamy, & Deouell, 2014; Võ & Wolfe, 2013). More similarly to our study, others have investigated the effects of syntactic consistencies (i.e., whether the object is placed in its typical within-scene location), and found comparably late effects, most prominently between 400 and 600 ms (Demiral, Malcolm, & Henderson, 2012; Võ & Wolfe, 2013). Such effects of syntactic consistencies have been linked to efficient behavioral performance in complex multi-object scenes (Draschkow & Võ, 2017).

How can the early effect observed here be reconciled with the late effects found for typical object placement within a scene? Studies on object-scene consistencies differ in two important ways from our study. First, in these studies participants are cued to look at the object’s location within the scene, whereby retinotopic object location is purposefully eliminated. Second, these studies measure overall waveform changes between congruent and incongruent conditions, which may reflect more general, post-perceptual signatures of consistency processing or object-scene interaction. By contrast, our analysis used highly sensitive decoding methods to explicitly focus on object discriminability within typically and atypically positioned objects, so that our results are best understood as a rapid signature of experience-based tuning of the visual architecture. Our findings and the findings of these previous studies thus highlight different facets of the processing of regularly structured environments, where both the early effects of typical retinotopic locations and the later effects of typical within-scene positioning may contribute to efficient real-world perception.

### 4.8 Conclusion

Together, our results demonstrate a general principle of object coding in human visual cortex: Information about an object’s location and its identity are processed interactively, where objects appearing in their typical retinotopic locations are more efficiently coded than objects appearing in atypical retinotopic locations. This processing advantage manifests in object decoding 140 ms after stimulus onset, suggesting that early object representations are tied to the object’s extensively experienced visual-field location. This finding can provide a novel explanation for the efficient parsing of complex real-world environments, which contain large numbers of typically positioned objects.

## Acknowledgements

The research was supported by a DFG Emmy-Noether Grant awarded to R.M.C. (CI241-1/1).

## References

Baeck, A., Wagemans, J., & Op de Beeck, H. P. (2013). The distributed representation of random and meaningful object pairs in human occipitotemporal cortex: the weighted average as a general rule. Neuroimage, 70, 37–47.

Baldassi, C., Alemi-Neissi, A., Pagan, M., DiCarlo, J. J., Zecchina, R., & Zoccolan, D. (2013). Shape similarity, better than semantic membership, accounts for the structure of visual object representations in a population of monkey inferotemporal neurons. PLoS Computational Biology, 9, e1003167.

Bar, M. (2004). Visual objects in context. Nature Reviews Neuroscience, 5, 617–629.

Bar, M., Kassam, K. S., Ghuman, A. S., Boshyan, J., Schmid, A. M., Dale, A. M., Hämäläinen, M. S., Marinkovic, K., Schacter, D. L., Rosen, B. R., & Halgren, E. (2006). Top-down facilitation of visual recognition. Proceedings of the National Academy of Sciences, U.S.A., 103, 449–454.

Biederman, I., Mezzanotte, R. J., & Rabinowitz, J. C. (1982). Scene perception: detecting and judging objects undergoing relational violations. Cognitive Psychology, 14, 143–177.

Bracci, S., Ritchie, J. B., & op de Beeck, H. P. (2017). On the partnership between neural representations of object categories and visual features in the ventral visual pathway. Neuropsychologia, 105, 153–164.

Brainard, D. H. (1997). The psychophysics toolbox. Spatial Vision, 10, 433–436.

Brandman, T., & Peelen, M. V. (2017). Interaction between scene and object processing revealed by human fMRI and MEG decoding. Journal of Neuroscience, 37, 7700–7710.

Carlson, T. A., Hogendoorn, H., Kanai, R., Mesik, J., & Turret, J. (2011). High temporal resolution decoding of object position and category. Journal of Vision, 11, 9.

Carlson, T. A., Tovar D. A., Alink, A., & Kriegeskorte, N. (2013). Representational dynamics of object vision: the first 1000 ms. Journal of Vision, 13, 1–19.

Chan, A. W., Kravitz, D. J., Truong, S., Arizpe, J., & Baker, C.I. (2010). Cortical representations of bodies and faces are strongest in commonly experienced configurations. Nature Neuroscience, 13, 417–418.

Chun, M. M. (2000). Contextual cueing of visual attention. Trends in Cognitive Sciences, 4, 170–178.

Cichy, R. M., Chen, Y., & Haynes, J. D. (2011). Encoding the identity and location of objects in human LOC. Neuroimage, 54, 2297–2307.

Cichy, R. M., Pantazis, D., & Oliva, A. (2014). Resolving human object recognition in space and time. Nature Neuroscience, 17, 455–462.

Cichy, R. M., Pantazis, D., & Oliva, A. (2016). Similarity-based fusion of MEG and fMRI reveals spatio-temporal dynamics in human cortex during visual object recognition. Cerebral Cortex, 26, 3563–3579.

Contini, E. W., Wardle, S. G., & Carlson, T. A. (2017). Decoding the time-course of object recognition in the human brain: From visual features to categorical decisions. Neuropsychologia, 105, 165–176.

de Haas, B., Schwarzkopf, D. S., Alvarez, I., Lawson, R. P., Henriksson, L., Kriegeskorte, N., & Rees, G. (2016). Perception and processing of faces in the human brain is tuned to typical facial feature locations. Journal of Neuroscience, 36, 9289–9302.

Demiral, S. B., Malcolm, G. L., & Henderson, J. M. (2012). ERP correlates of spatially incongruent object identification during scene viewing: contextual expectancy versus simultaneous processing. Neuropsychologia, 50, 1271–1285.

Draschkow, D., & Võ, M. L.-H. (2017). Scene grammar shapes the way we interact with objects, strengthens memories, and speeds search. Scientific Reports, 7, 16471.

Fahrenfort, J. J., van Leeuwen, J., Olivers, C. N. L., & Hogendoorn, H. (2017). Perceptual integration without conscious access. Proceedings of the National Academy of Sciences, U.S.A., 114, 3744–3749.

Ganis, G., & Kutas, M. (2003). An electrophysiological study of scene effects on object identification. Cognitive Brain Research, 16, 123–144.

Golomb, J. D., & Kanwisher, N. (2012). Higher level visual cortex represents retinotopic, not spatiotopic, object location. Cerebral Cortex, 22, 2794–2810.

Gronau, N., & Shachar, M. (2014). Contextual integration of visual objects necessitates attention. Attention, Perception & Psychophysics, 76, 695–714.

Hasson, U., Levy, I., Behrmann, M., Hendler, T., & Malach, R. (2002). Eccentricity bias as an organizing principle for human high-order object areas. Neuron, 34, 479–490.

Hayhoe, M., & Ballard, D. (2005). Eye movements in natural behavior. Trends in Cognitive Sciences, 9, 188–194.

Hemond, C. C., Kanwisher, N., & Op de Beeck, H. P. (2007). A preference for contralateral stimuli in human object- and face-selective cortex. PLoS One, 2, e574.

Henriksson, L., Mur, M., & Kriegeskorte, N. (2015). Faciotopy – A face-feature like topology in the human occipital face area. Cortex, 72, 156–167.

Hong, H, Yamins, D. L. K., Majaj, N. J., & DiCarlo, J. J. (2016). Explicit information for category-orthogonal object properties increases along the ventral visual stream. Nature Neuroscience, 19, 613–622.

Isik, L., Meyers, E. M., Leibo, J. Z., & Poggio, T. (2014). The dynamics of invariant object recognition in the human visual system. Journal of Neurophysiology, 111, 91–102.

Issa, E. B., & DiCarlo, J. J. (2012). Precedence of the eye region in neural processing of faces. Journal of Neuroscience, 32, 16666–16682.

Kaiser, D., Azzalini, D. C., & Peelen, M. V. (2016). Shape-independent object category responses revealed by MEG and fMRI decoding. Journal of Neurophysiology, 115, 2246–2250.

Kaiser, D., & Haselhuhn, T. (2017). Facing a regular world: How spatial object structure shapes visual processing. Journal of Neuroscience, 37, 1965–1967.

Kaiser, D., Oosterhof, N. N., & Peelen, M. V. (2016). The neural dynamics of attentional selection in natural scenes. Journal of Neuroscience, 36, 10522–10528.

Kaiser. D., & Peelen, M. V. (2018). Transformation from independent to integrative coding of multi-object arrangements in human visual cortex. Neuroimage, 169, 334–341.

Kaiser, D., Stein, T., & Peelen, M. V. (2014). Object grouping based on real-world regularities facilitates perception by reducing competitive interactions in visual cortex. Proceedings of the National Academy of Sciences, U.S.A., 111, 11217–11222.

Kaiser, D., Stein, T., & Peelen, M. V. (2015). Real-world spatial regularities affect visual working memory for objects. Psychonomic Bulletin & Review, 22, 1784–1790.

Kravitz, D. J., Kriegeskorte, N., & Baker, C. I. (2010). High-level visual object representations are constrained by position. Cerebral Cortex, 20, 2916–2925.

Kravitz, D. J., Saleem, K. S., Baker, C. I., Ungerleider, L. G., & Mishkin, M. (2013). The ventral visual pathway: an expanded neural framework for the processing of object quality. Trends in Cognitive Sciences, 17, 26–49.

Kravitz, D. J., Vinson, L. D., & Baker, C. I. (2008). How position dependent is visual object recognition? Trends in Cognitive Sciences, 12, 114–122.

Land, M. F. (2009). Vision, eye movements, and natural behavior. Visual Neuroscience, 26, 51–62.

Mudrik, L., Lamy, D., & Deouell, L. Y. (2010). ERP evidence for context congruity effects during simultaneous object-scene processing. Neuropsychologia, 48, 507–517.

Mudrik, L., Shalgi, L., Lamy, D., & Deouell, L. Y. (2010). ERP correlates of spatially incongruent object identification during scene viewing: contextual expectancy versus simultaneous processing. Neuropsychologia, 56, 447–458.

Oliva, A., & Torralba, A. (2007). The role of context in object recognition. Trends in Cognitive Sciences, 11, 520–527.

Oostenveld, R., Fries, P., Maris, E., & Schoffelen, J. M. (2011). Fieldtrip: open source software for advances analysis of MEG, EEG, and invasive electrophysiological data. Computational Intelligence and Neuroscience, 2011, 156869.

Oosterhof, N. N., Connolly, A. C., & Haxby, J. V. (2016). CoSMoMVPA: multi-modal multivariate pattern analysis of neuroimaging data in Matlab / GNU Octave. Frontiers in Neuroinformatics, 10, 20.

Peelen, M. V., & Downing, P. E. (2017) Category selectivity in human visual cortex: Beyond visual object recognition. Neuropsychologia, 105, 177–183.

Poldrack, R. A. (2007). Region of interest analysis for fMRI. Social Cognitive and Affective Neuroscience, 2, 67–70.

Rouder, J. N., Speckman, P. L., Sun, D., Morey, R. D., & Iverson, G. (2009). Bayesian t-tests for accepting and rejecting the null hypothesis. Psychonomic Bulletin & Review, 16, 225–237.

Russell, B. C., Torralba, A., Murphy, K. P., & Freeman, W. T. (2008). LabelMe: a database and web-based tool for image annotation. International Journal of Computer Vision, 77, 157–173.

Schwarzlose, R. F., Swisher, J. D., Dang, S., & Kanwisher, N. (2008). The distribution of category and location information across object-selective regions in human visual cortex. Proceedings of the National Academy of Sciences, U.S.A., 105, 4447–4452.

Smith, S. M., & Nichols, T. E. (2009). Threshold-free cluster enhancement: addressing problems of smoothing, threshold dependence and localization in cluster interference. Neuroimage, 66, 215–222.

Stein, T., Kaiser, D., & Peelen, M. V. (2015). Interobject grouping facilitates visual awareness. Journal of Vision, 15, 10.

Turano, K. A., Geruschat, D. R., & Baker, F. H. (2003). Oculomotor strategies for the direction of gaze tested with a real-world activity. Vision Research, 43, 333–346.

Vinckier, F., Dehaene, S., Jobert, A., Dubus, J. P., Sigman, M., & Cohen, L. (2007). Hierarchical coding of letter strings in the ventral stream: dissecting the inner organization of the visual word-form system. Neuron, 55, 143–156.

Võ, M. L.-H., & Wolfe, J. M. (2013). Differential ERP signatures elicited by semantic and syntactic processing in scenes. Psychological Science, 24, 1816–1823.

Willenbockel, V., Sadr, J., Fiset, D., Horne, G. O., Gosselin, F., & Tanaka, J. W. (2010). Controlling low-level image properties: The SHINE toolbox. Behavior Research Methods, 42, 671–684.

Wolfe, J. M., Võ, M. L.-H., Evans, K. K., & Greene, M. R. (2011). Visual search in scenes involves selective and nonselective pathways. Trends in Cognitive Sciences, 15, 77–84.

